# A novel MRI-based data fusion methodology for efficient, personalised, compliant simulations of aortic haemodynamics

**DOI:** 10.1101/2021.05.15.444156

**Authors:** Catriona Stokes, Mirko Bonfanti, Zeyan Li, Jiang Xiong, Duanduan Chen, Stavroula Balabani, Vanessa Díaz-Zuccarini

## Abstract

We present a novel, cost-efficient methodology to simulate aortic haemodynamics in a patient-specific, compliant aorta using an MRI data fusion process. Based on a previously-developed Moving Boundary Method, this technique circumvents the high computational cost and numerous structural modelling assumptions required by traditional Fluid-Structure Interaction techniques. Without the need for Computed Tomography (CT) data, the MRI images required to construct the simulation can be obtained during a single imaging session. Black Blood MR Angiography and 2D Cine-MRI data were used to reconstruct the luminal geometry and calibrate wall movement specifically to each region of the aorta. 4D-Flow MRI and non-invasive pressure measurements informed patient-specific inlet and outlet boundary conditions. Luminal area closely matched 2D Cine-MRI measurements with a mean error of less than 4.6% across the cardiac cycle, while physiological pressure and flow distributions were simulated to within 3.3% of patient-specific targets. Moderate agreement with 4D-Flow MRI velocity data was observed. Despite lower peak velocity, an equivalent rigid-wall simulation predicted a mean Time-Averaged Wall Shear Stress (TAWSS) 13% higher than the compliant simulation. The agreement observed between compliant simulation results and MRI data is testament to the accuracy and efficiency of this MRI-based simulation technique.

## 1. Introduction

Computational Fluid Dynamics (CFD) and phase-contrast Magnetic Resonance Imaging (PC-MRI) techniques such as Four-Dimensional Flow MRI (4DMR) facilitate the analysis of arterial haemodynamics, providing valuable insights to support clinical decision-making. Due to limitations in spatiotemporal resolution, 4DMR cannot accurately capture small-scale flow features such as the fluid boundary layer. It hence cannot accurately estimate clinically relevant indices such as Wall Shear Stress (WSS) that are implicated in the onset and development of various cardiovascular diseases (Castagna et al., 2021; Mazzi et al., 2020; Miyazaki et al., 2017; Piatti et al., 2017; Zimmermann et al., 2018). Informing CFD simulations with medical imaging data can facilitate both high resolution and patient-specific accuracy, yielding higher-quality haemodynamic data than any individual modality could provide. However, simulation accuracy depends on the choice of modelling assumptions, such as vessel wall compliance.

Modelling wall compliance is fraught with difficulties, so a rigid-wall assumption is commonly used. Unfortunately, this assumption has been shown to significantly affect the accuracy of haemodynamic metrics such as WSS (Lantz et al., 2011; Miyazaki et al., 2017; Qiao et al., 2019). Fluid-Structure Interaction (FSI), a technique that couples the structural dynamics of the vessel wall with the flow solution, is traditionally used to simulate compliance. However, the complex, inhomogeneous material properties of the aortic wall cannot be directly measured *in-vivo*. Instead, a constant literature value of Young’s Modulus (*E*) is often assumed, which fails to accurately capture vessel wall movement throughout the aorta (He et al., 2021; Ryzhakov et al., 2019; Saitta et al., 2019; Tang et al., 2020).

Bonfanti et al. (2017) developed a Moving Boundary Method (MBM) to circumvent the structural assumptions and computational cost associated with FSI. Using CT data to reconstruct the aortic lumen and 2D Cine-MRI (Cine-MRI) data to compute vessel wall compliance locally in each region of the aorta, the MBM can accurately capture wall movement throughout the aorta, unlike FSI. MBM simulations of Type-B Aortic Dissection (TBAD), a severe pathology characterised by a tear in the innermost layer of the aortic wall, agreed closely with an equivalent FSI simulation yet required only half the simulation time (Bonfanti et al., 2018). However, CT images are not always available, for example, in healthy patients where the exposure to high doses of ionising radiation is not clinically justified. MRI-based techniques have been developed but typically employ a rigid-wall assumption (Bozzi et al., 2017; Madhavan and Kemmerling, 2018; Youssefi et al., 2017).

Several MRI-based FSI studies of healthy aortae have appeared in the literature, including those of Lantz et al. (2011), Boccadifuoco et al. (2018) and Pons et al. (2020). Each used a constant value of *E* throughout the aorta, requiring multiple FSI simulations for its calibration. Lantz et al. (2011) and Boccadifuoco et al. (2018) used literature values of *E* in the physiological range, whereas Pons et al. (2020) iteratively adjusted *E* by calibrating the pulse wave velocity (PWV) using estimations from 4DMR. When compared against MRI data, FSI failed to accurately capture wall movement throughout the aorta with a uniform *E*. Thus, there remains a need for cost-efficient, MRI-based CFD methods to accurately characterise non-uniform arterial compliance.

This study presents a novel MBM-based methodology to construct patient-specific, compliant simulations of a healthy thoracic aorta using a unique MRI data fusion process. MRA and Cine-MRI are used to segment the aorta and calibrate region-specific wall movement, while 4DMR and non-invasive pressure measurements inform patient-specific boundary conditions. Compared with FSI, the proposed CFD methodology can provide accurate results in significantly less time, using data from a single MRI acquisition session. Thus, this technique could facilitate shorter clinical decision-making timescales and reduce clinical resource requirements.

## 2. Methods

### 2.1. Clinical Data

Three MRI sequences of the thoracic aorta of a healthy volunteer were acquired using a Philips Achieva 3.0 TX multi-source MRI scanner at Beijing Institute of Technology, Beijing, China (Philips Medical Systems, Holland). MRA images of the thoracic aorta at diastole were acquired with a resolution of 0.48 × 0.48 × 1 *mm*^3^. Cine-MRI images were acquired as transverse planes with a resolution of 1.25 × 1.25 × 4 *mm*^3^ and a timestep of 38 ms. Finally, sagittal 4DMR images were acquired at 24 points across the cardiac cycle with a voxel size of 2.5 × 2.5 × 2.8 *mm*^3^ and a single velocity encoding (VENC) of 1 m/s. A heart rate of 68 beats per minute and systolic and diastolic brachial blood pressures (*P*_*sb*_, *P*_*db*_) of 117 mmHg and 72 mmHg were measured using a sphygmomanometer after MRI acquisition. This work was ethically approved by the Institutional Review Board of the Chinese PLA General Hospital (S201703601).

### 2.2. Geometry and Meshing

The simulation workflow is shown schematically in Figure 1. A patient-specific luminal geometry was reconstructed by manual segmentation of MRA images using Mimics (Materialize NV, Leuven, Belgium) (Figure 1a). This diastolic segmentation was bounded by an inlet at the ascending aorta (AA) and an outlet at the abdominal aorta (AbAo). Proximal sections of the supra-aortic branches, the Brachiocephalic Trunk (BT), Left Common Carotid (LCC), and Left Subclavian Artery (LSA), were included, with outlets at the terminus of each. As MRA could not resolve the intercostal arteries, they were not included in the segmentation. As such, flow loss through these branches, measured as 5-10% of the stroke volume (SV), was not modelled.

**Figure 1:**
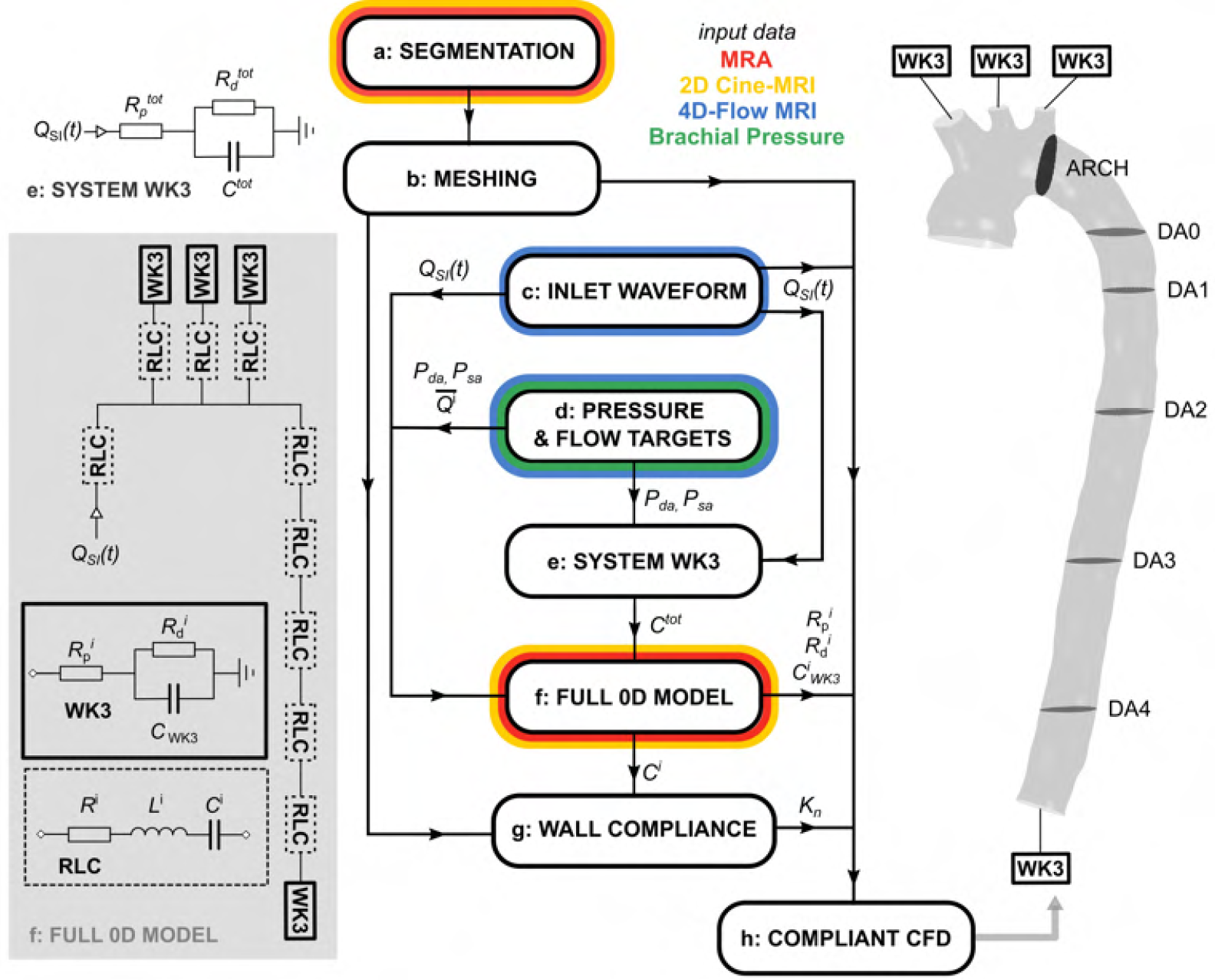
Flowchart of the simulation methodology including schematic diagrams of the system WK3 (e), the 0D domain (f) and the 3D CFD domain (h). Steps (a) through (h) are referred to throughout Section 2, where they are each described in detail.

Cross-sectional area measurements in the segmented proximal aorta were approximately 30% smaller than the Cine-MRI areas at diastole in the same locations. This is likely due to separation-induced flow stagnation in this region of the MRA images, which can reduce image contrast and erroneously indicate the presence of wall tissue (Henningsson et al., 2020). Fusing MRA and Cine-MRI, selected regions of the segmented aorta were dilated by up to 2 mm in the surface-normal direction using Simpleware ScanIP (Synopsis Inc., CA, USA) to match the diastolic (minimum) cross-sectional area measurements from Cine-MRI whilst preserving the morphology of the lumen. The segmented geometry was used to construct a tetrahedral computational mesh with 525k elements using Ansys Fluent 20.0 (Ansys Inc., PA, USA) (Figure 1b). Details on prismatic layer settings are provided in Appendix A. Mesh refinement was determined using a mesh independence study described in Appendix B.

### 2.3. Boundary Conditions

The choice of inlet and outlet boundary conditions are critical in accurately simulating patient-specific physiological flow and pressure distributions. The patient-specific inlet flow rate waveform was extracted from 4DMR data at the ascending aorta using GTFlow (GyroTools LLC., Zurich, Switzerland). Using spline interpolation in MATLAB (MathWorks Inc., Natick, Massachusetts, USA), a smooth waveform with 1 ms increments, *Q*_*SI*_(*t*), was derived from these measurements and used to apply a uniform inlet velocity profile (IVP) *v*_*inlet*_(*t*) = *Q*_*SI*_(*t*)/*A*_*inlet*_ (Figure 1c). Compared to a patient-specific, three-component (3D) IVP extracted from 4DMR, the choice of a uniform IVP is likely to impact the accuracy of velocity and WSS distributions in the ascending aorta and aortic arch. It is expected to have a lesser impact on flow in the descending aorta (DA) according to Armour et al. (2021); Pirola et al. (2018), though the uniform IVP may under-predict flow helicity and radial velocity, particularly during late systolic deceleration (Morbiducci et al., 2013; Youssefi et al., 2017). Applying a non-uniform measured IVP may lead to improved velocity agreement between CFD and 4DMR; however, doing so is non-trivial. Adding this step would not affect any other stages of the proposed compliance modelling methodology, so a uniform IVP was deemed suitable for this study. The inlet was fixed in place and not permitted to dilate across the cardiac cycle.

Three-element Windkessel (WK3) outlet pressure boundary conditions were applied at each CFD outlet to simulate the effects of the peripheral vascular system and minimise the incidence of non-physical pressure wave reflections. Each WK3 is an electrical analogue consisting of a proximal resistance *R*_*p*_, a distal resistance *R*_*d*_ and a capacitance *C*_*WK*3_, whose values must be carefully calibrated to the specific patient.

WK3 calibration began by determining target values of aortic systolic and diastolic pressure (*P*_*sa*_, *P*_*da*_) and mean flow rates at each outlet (Figure 1d). Diastolic pressure remains relatively constant throughout the arterial tree, so *P*_*da*_ was set to *P*_*db*_. *P*_*sa*_ was derived from *P*_*sb*_ using *P*_*sa*_ ≈ 0.83*P*_*sb*_ + 0.15*P*_*db*_ (Westerhof et al., 2010). The target mean flow rate at each outlet was then derived from 4DMR data. As one voxel accounted for 10-30% of the cross-sectional area of each supra-aortic branch, measurements of flow rate at these locations incurred high uncertainties. Instead, flow rate waveforms were first extracted at four planes in the DA (DA1 - DA4 in Fig. 1). The total mean flow rate through the supra-aortic branches, 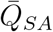, was then calculated as

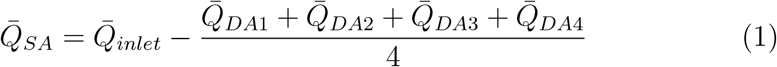

and 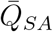 was split proportionally between BT, LCC and LSA based on their diastolic cross-sectional areas.

Next, a single-WK3 analogue of the full arterial system (Figure 1e) was used to determine the total arterial compliance, *C*^*tot*^, with a method described by Les et al. (2010). Using *Q*_*SI*_ (*t*) at the inlet of the WK3, 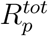, 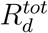, and *C*^*tot*^ were adjusted iteratively until target values of *P*_*sa*_ and *P*_*da*_ were observed at the inlet.

A full 0D lumped parameter model of the aorta was then constructed (Figure 1f). In the full 0D model, the CFD domain was represented by nine discrete *RLC* units, each containing a resistor (*R*^*i*^), an inductor (*L*^*i*^) and a capacitor 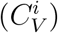 to simulate fluid pressure loss, inertance and volume compliance due to wall movement, respectively. The inlet flow rate waveform *Q*_*SI*_(*t*) was applied at the inlet, and a WK3 was connected to each outlet, each with parameters 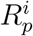, 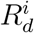, and 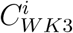.

To compute 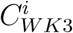, the peripheral compliance 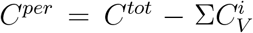 was 153 divided between each outlet proportionally to the mean flow rate of each supra-aortic branch, 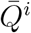:

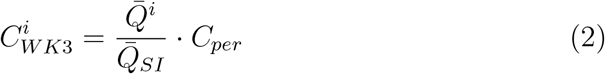

The total resistance of each outlet was calculated using:

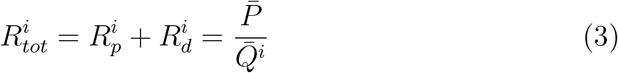

where 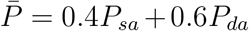. An initial guess of 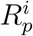 was obtained to match the characteristic impedance of the vessel:

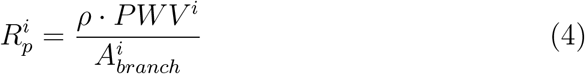

where 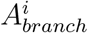 is the cross-sectional area of the branch outlet.

For each *RLC* unit, *R*^*i*^ were computed using the total pressure loss through each segment from a steady-state CFD simulation at the mean inlet flow rate. *L*^*i*^ were calculated using an expression for large arteries of length *l*^*i*^ and cross-sectional area *A*^*i*^:

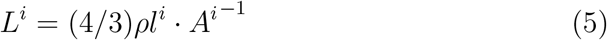

where *ρ* is the density of blood (Westerhof et al., 2010). The volume compliances 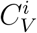 were calculated as

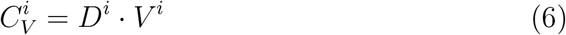

where *V*^*i*^ is the volume of the segment. *D*^*i*^ is the local area distensibility which is calculated from Cine-MRI measurements of cross-sectional area at each segment as

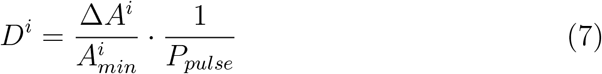

where Δ*A*^*i*^ is the maximum cross-sectional area change across the cardiac cycle and 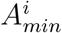 is the diastolic area. Because the supra-aortic branches were not resolved by Cine-MRI, their respective area distensibility values were calculated using an empirical relationship from Reymond et al. (2009):

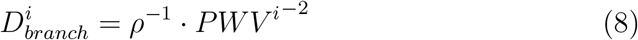

where 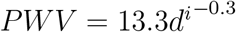 and *d*^*i*^ is the branch diameter.

The full 0D model results in a system of ordinary differential equations which were solved numerically using 20-sim (ControllabProducts B.V., Enschede, The Netherlands) by backward differentiation. 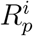 values were manipulated from their initial estimate until target values of *P*_*sa*_ and *P*_*da*_ were observed at the inlet. The final WK3 parameters are shown in Table 1.

**Table 1:**
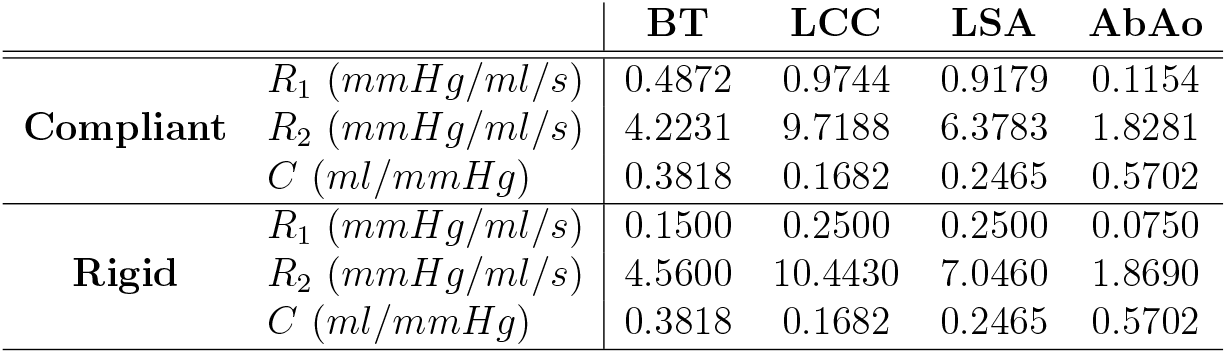
Patient-specific three-element Windkessel parameters for compliant and rigid CFD simulations, determined using the 0D tuning process described in Section 2

### 2.4. Wall Compliance

In our moving mesh approach (Bonfanti et al., 2017), the structural dynamics of the wall are not explicitly modelled. Instead, the magnitude of displacement of each mesh node *n* at the aortic wall, *δ*_*n*_, is computed as a linear function of the local pressure in the surface-normal direction **n**_*n*_:

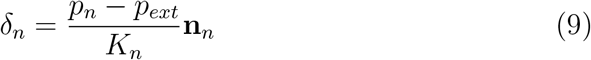

where *K*_*n*_ is the local stiffness coefficient, *p*_*n*_ is the fluid pressure, and *p*_*ext*_ is the pressure exerted from the external side of the aortic wall, set as *P*_*da*_. *K*_*n*_ is calculated (Figure 1g) using:

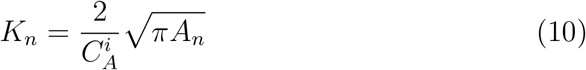

where *A*_*n*_ is the cross-sectional area of the lumen at node *n*. 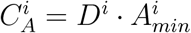 is the area compliance in the region of node *n*, where *D*^*i*^ values are obtained from equation 7. As the inlet and outlets are fixed in place and do not dilate, the faces bordering them were given an artificially high stiffness to prevent excessive cell distortion. Finally, five smoothing operations were performed to eliminate the discontinuities in stiffness between each aortic region.

We also performed a simulation with rigid walls to assess the impact of compliance on parameters of interest. In the rigid 0D model, the capacitors in each *RLC* unit representing the CFD domain were removed, and identical pressure and flow rate values were targeted. This resulted in a separate set of WK3 parameters for each simulation (Table 1).

### 2.5. CFD Simulation

The transient, three-dimensional Navier-Stokes equations were solved numerically using the finite-volume solver Ansys CFX 20.0 (Figure 1h). Blood was modelled as an incompressible non-Newtonian fluid using the CarreauYasuda viscosity model with empirical constants from Gijsen et al. (1999) and a density of 1056 *kg/m*^3^. Using the Reynolds number (*Re*) definitions for pulsatile cardiovascular flow from Peacock et al. (1998), a nominal shear rate defined by Cagney and Balabani (2019), and the peak velocity from 4DMR, the peak *Re*_*p*_ of 4157 was observed to exceed the critical *Re*_*c*_ of 3505, indicating the onset of turbulence. The k-*ω* Shear Stress Transport (SST) Reynolds-Averaged formulation of the Navier-Stokes equations was employed to model turbulence due to its ability to predict the onset and amount of flow separation under adverse pressure gradients (Lantz et al., 2011). A low turbulence intensity of 1% was applied at the inlet and outlets, which was found to most accurately represent healthy aortic flow by Kousera et al. (2013). Timesteps of 1 ms were solved using the implicit, second-order backward-Euler method. A root-mean-square residual value of 10^−5^ was achieved for all equations within each timestep during the final cycle.

Simulations were run until periodic conditions were achieved, defined as less than 1% change in systolic and diastolic pressures between cycles, requiring five cycles in the compliant simulation (CS) and three in the rigid simulation (RS). The RS and CS required 2.5 and 9.2 hours per cycle on 8 Intel Xeon cores, respectively. Based on a direct comparison by (Bonfanti et al., 2018) on the same workstation, an equivalent FSI simulation would require approximately double the simulation time as the MBM simulations (≈ 18h per cycle).

## 3. Results

Inlet pressure and mean outlet flow rates were compared with target values to assess whether the WK3 BCs yielded physiological pressure and flow conditions. Targets for *P*_*sys*_, *P*_*dia*_ and 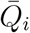 were achieved within 3.3% for both RS and CS (Table 2).

**Table 2:** Simulated compliant and rigid inlet systolic and diastolic pressure and mean outlet flow rates compared with target values. The percentage error between simulated and target values are indicated.

| Quantity | Measurement | Target | Rigid / % err | Compliant / % err |
| --- | --- | --- | --- | --- |
| $P_{sys}$ (mmHg) | 117 | 108 | 104.9 / 2.9% | 105.1 / 2.7% |
| $P_{dia}$ (mmHg) | 72 | 72 | 69.6 / 3.3% | 71.7 / 0.4% |
| $\bar{Q}_{BT}$ (ml/s) | 13.57 | 18.34 | 18.20 / 0.8% | 17.99 / 1.9% |
| $\bar{Q}_{LCC}$ (ml/s) | 6.17 | 8.11 | 8.02 / 1.1% | 7.99 / 1.5% |
| $\bar{Q}_{LSA}$ (ml/s) | 4.37 | 11.84 | 11.52 / 2.7% | 11.51 / 2.8% |
| $\bar{Q}_{AbAo}$ (ml/s) | 44.60 | 44.44 | 44.20 / 0.4% | 44.73 / 0.9% |

Simulated volume flow rate across the cardiac cycle was compared against 4DMR at DA1-DA4 (Figure 2). As intercostal flow loss was not simulated, every 4DMR point on each analysis plane (DA*) was linearly scaled by 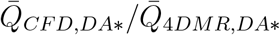 such that the SV in CFD and 4DMR were equal on plane DA*. This scaling facilitates a more direct comparison of the flow rate waveforms, even though measured flow loss is small (5-10%) (Baumler et al., 2020). Raw 4DMR measurements are also shown as a band encapsulating ±22ml of SV, representing the mean uncertainty for single-VENC 4DMR data (Kroeger et al., 2021). The peak flow rate is slightly over-predicted by the CS, an effect which is more pronounced in the proximal DA, and is under-predicted by the RS.

**Figure 2:**
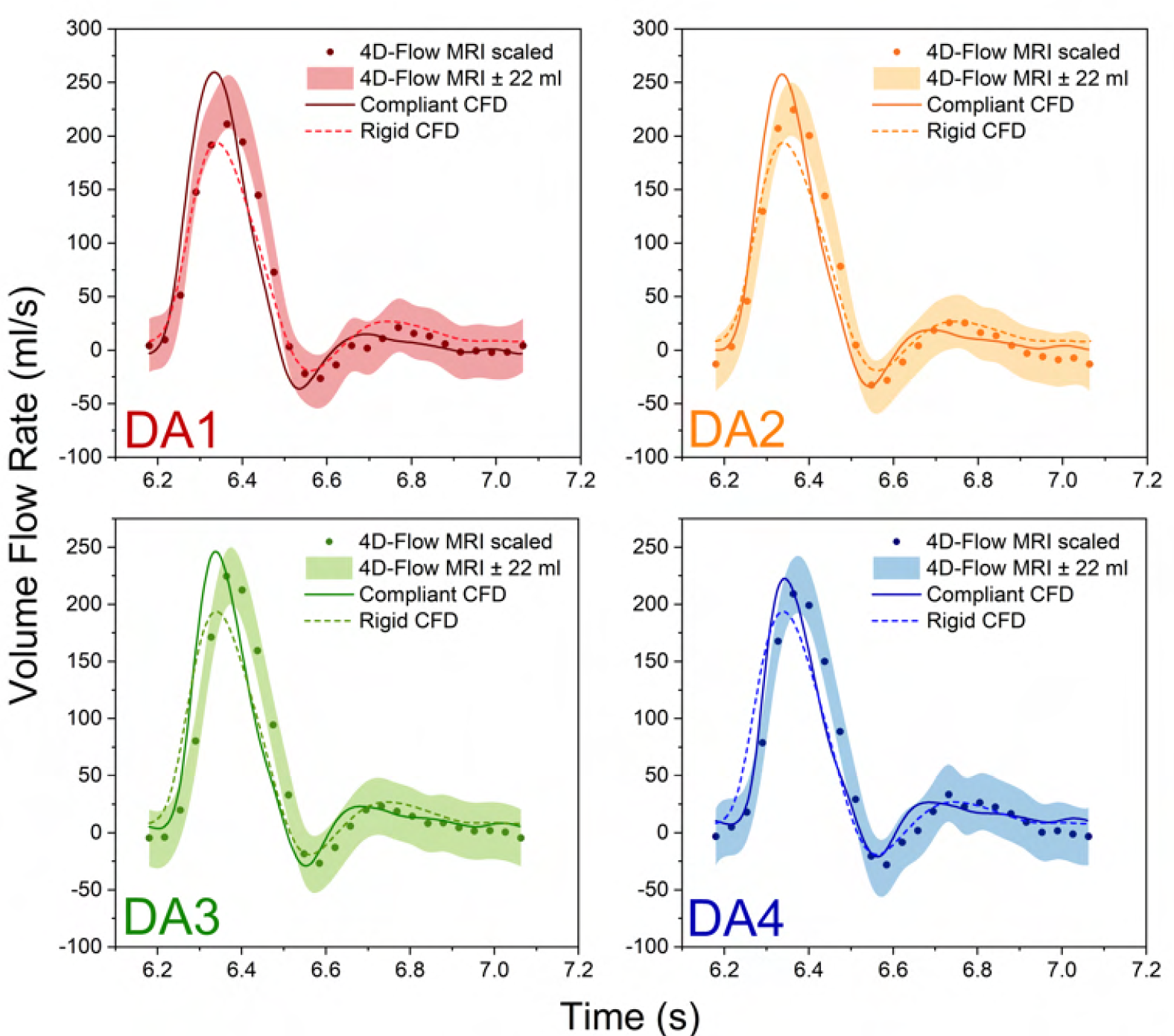
Volume flow rate comparison between CS, RS and 4DMR at four planes in the DA (see Figure 4 for locations). 4DMR results are shown as a band with a ± 22 ml uncertainty in SV, found to be the mean uncertainty for single-VENC 4DMR measurements of flow rate by Kroeger et al. (2021). 4DMR results are also shown as discrete measurements, with the SV scaled to match CFD due to the lack of 4DMR mass conservation.

Simulated luminal cross-sectional area changes across the cardiac cycle were compared with Cine-MRI measurements at the arch and DA1-DA4 (Figure 3). In addition to the measured diastolic area from Cine-MRI, the discrepancy in peak systolic area change between CFD and Cine-MRI and the mean error at each point on the waveform are indicated on each plot. Excellent agreement was observed throughout the DA, with less than 3% peak error and 2% mean error at all planes. Maximum and minimum areas matched closely at the arch, with less than 0.5% peak error. However, with an in-plane resolution of only 4 mm × 1.25 mm, Cine-MRI data at the arch appears noisy, and CFD and MRI waveforms do not appear as closely aligned, resulting in a 4.6% mean error. In addition, CFD yielded a faster rate of area expansion during the acceleration phase, an earlier peak, and a more pronounced secondary peak at end-systole compared to Cine-MRI.

**Figure 3:**
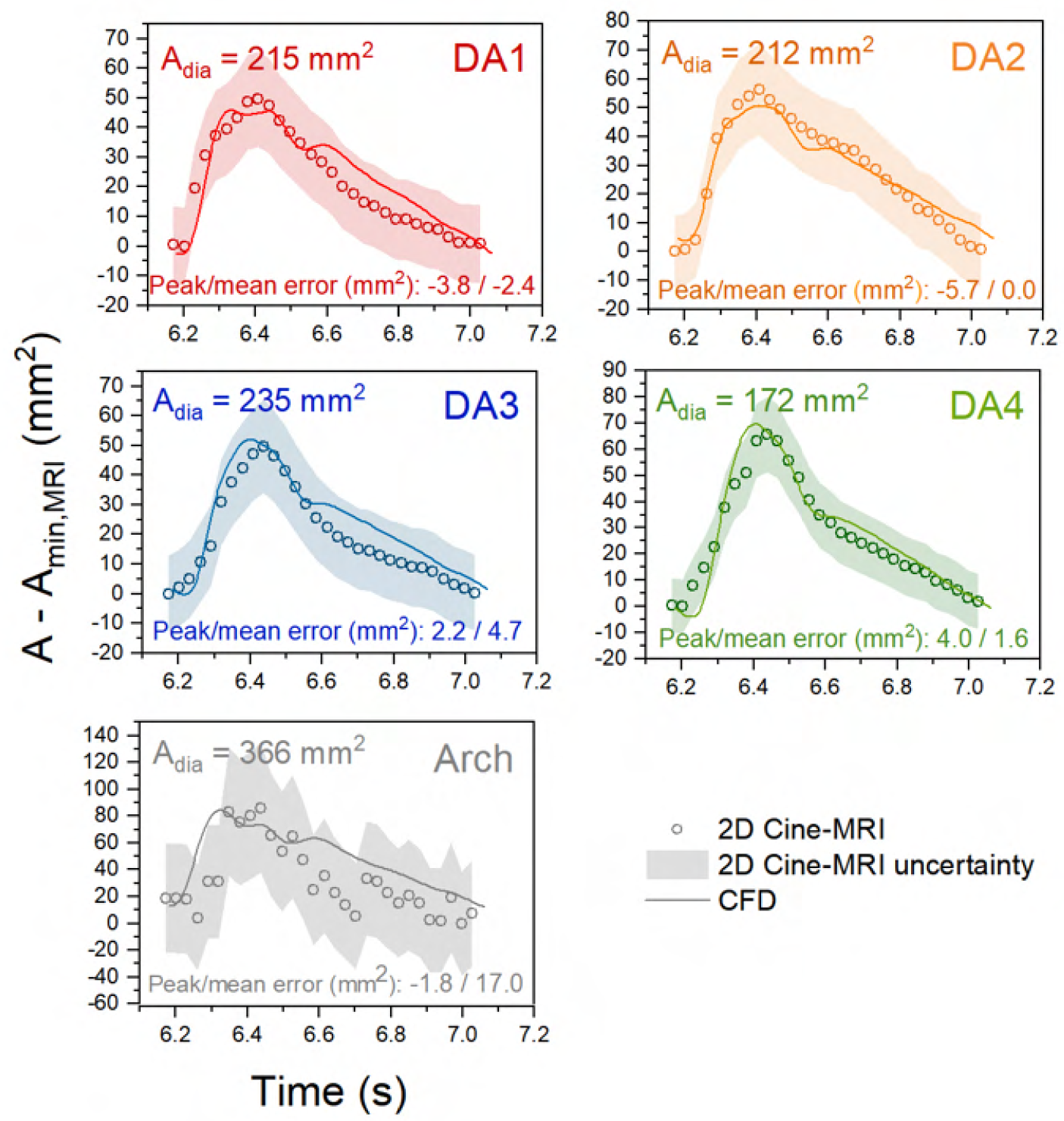
Luminal cross-sectional area change waveforms comparing Cine-MRI and compliant CFD at the arch and DA across the cardiac cycle. Absolute peak and mean errors are indicted along with the measured diastolic area from Cine-MRI. A band encapsulating ± 6% error are indicated in the DA planes and ± 12% at the arch. These errors were calculated by manually selecting the smallest and largest areas that could reasonably be chosen at each plane in GTFlow. See Figure 4 for plane locations.

Velocity magnitude contours from RS and CS were compared with 4DMR data at peak systole (T1, Figure 4) and late systole, as flow is decelerating (T2, Figure 5). At T2, CFD data was extracted such that the flow rate was equal to 4DMR at each plane. Note that CFD analysis planes were chosen to match the angle and location of 4DMR planes and thus were not perpendicular to the centreline.

**Figure 4:**
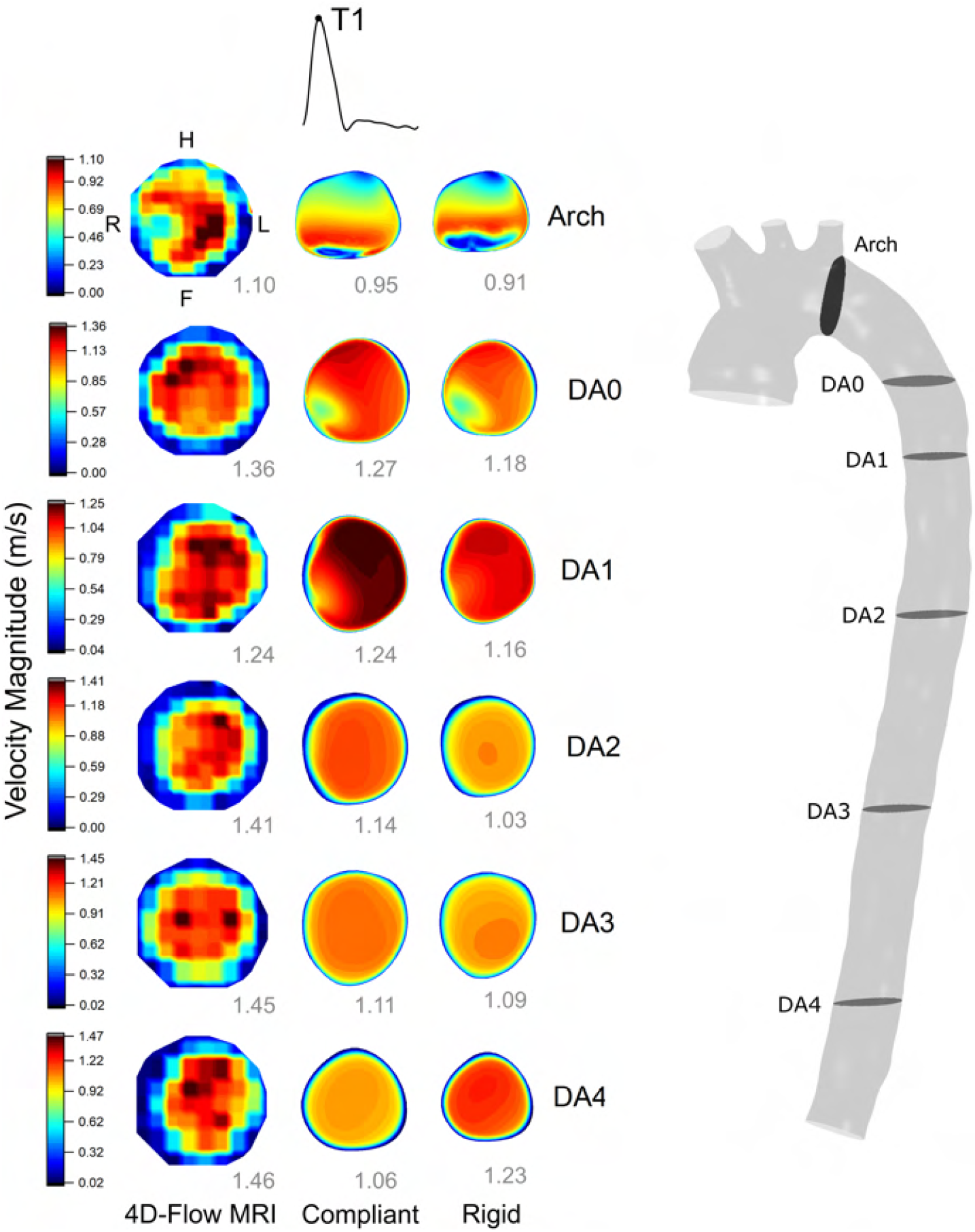
Velocity contour comparison between 4DMR, compliant and rigid CFD simulations at peak systole throughout the aorta. Velocity contour ranges are set by the minimum and maximum values from 4D Flow-MRI, and the peak velocity magnitude at each CFD plane is indicated at the bottom right of each contour. The arch plane orientation is indicated, while the orientation of the DA planes are indicated in Figure 5

**Figure 5:**
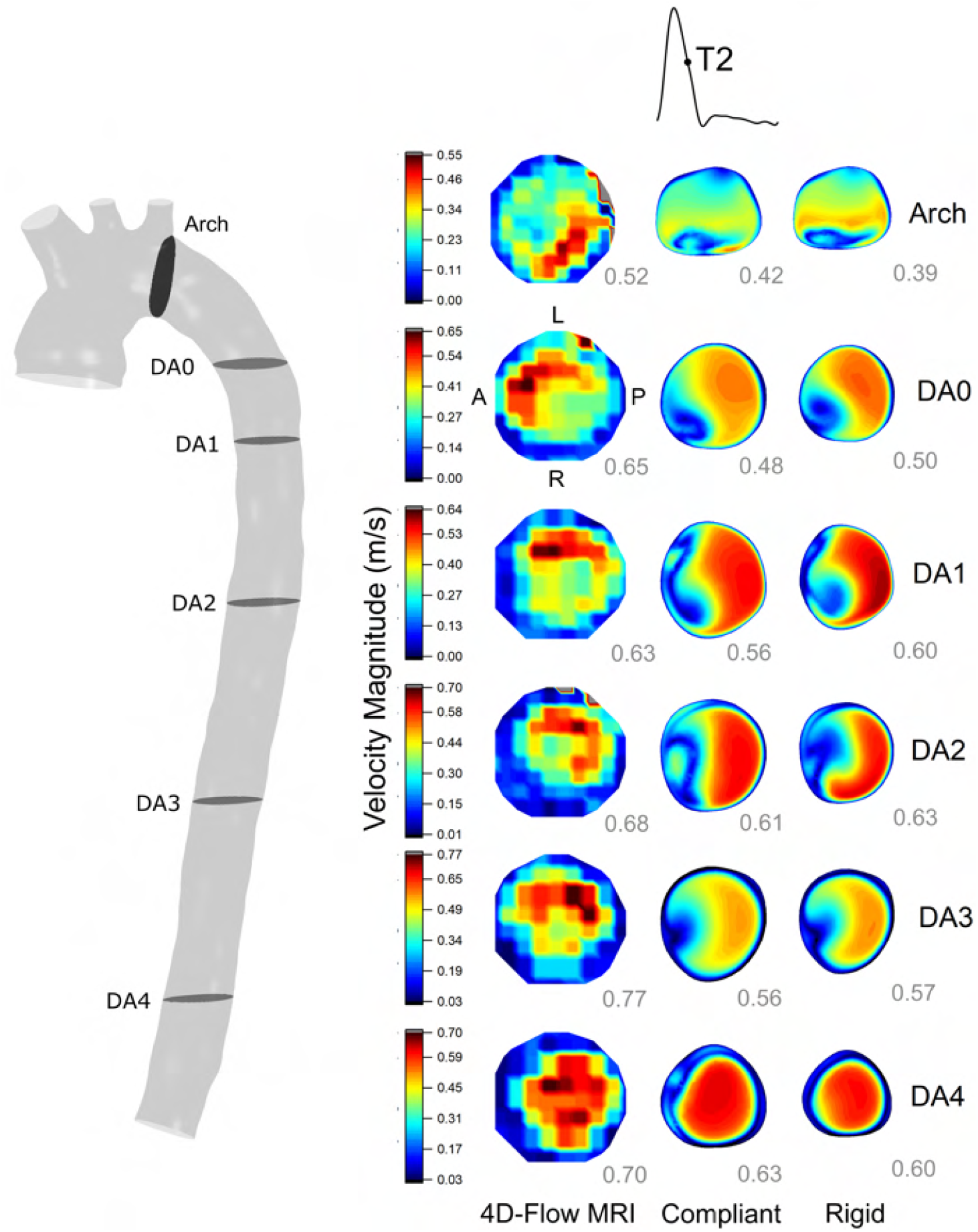
Velocity contour comparison between 4DMR, CS and RS at mid-deceleration throughout the aorta. Velocity contour ranges are set by the minimum and maximum values from 4D Flow-MRI, and the peak velocity magnitude at each CFD plane are indicated at the bottom right of each contour. The DA plane orientation is indicated, while the orientation of the arch planes are indicated on Figure 4

A moderate qualitative agreement in velocity distribution is observed between CFD and 4DMR. At the arch, MRI shows that flow is more concentrated to the left side than CFD. At T1, regions of high velocity in the DA correspond closely between MRI and CFD, particularly at DA0 and DA1. At T2, CFD predicts a high-velocity region that is concentrated towards the posterior wall, whereas in MRI, it progressively rotates from the anterior to the left posterior wall along the DA.

Peak velocity tends to increase along the DA in both CFD and 4DMR. It is almost universally under-predicted by CFD, by 0-28% at T1 and 10-28% at T2. The agreement is closest at T1 in the proximal aorta, where the compliant peak flow rate is over-predicted. Flow rates are precisely matched between CFD and MRI at each plane at T2, where under-prediction in peak velocity is universal. The reasons for the observed under-prediction will be discussed in Section 4.

Using the definitions from Gallo et al. (2012), Time Averaged Wall Shear Stress (TAWSS) and Oscillatory Shear Index (OSI) contours from the CS are shown alongside contours of difference between CS and RS in Figure 6, with simulated velocity streamlines to assist in their interpretation.

**Figure 6:**
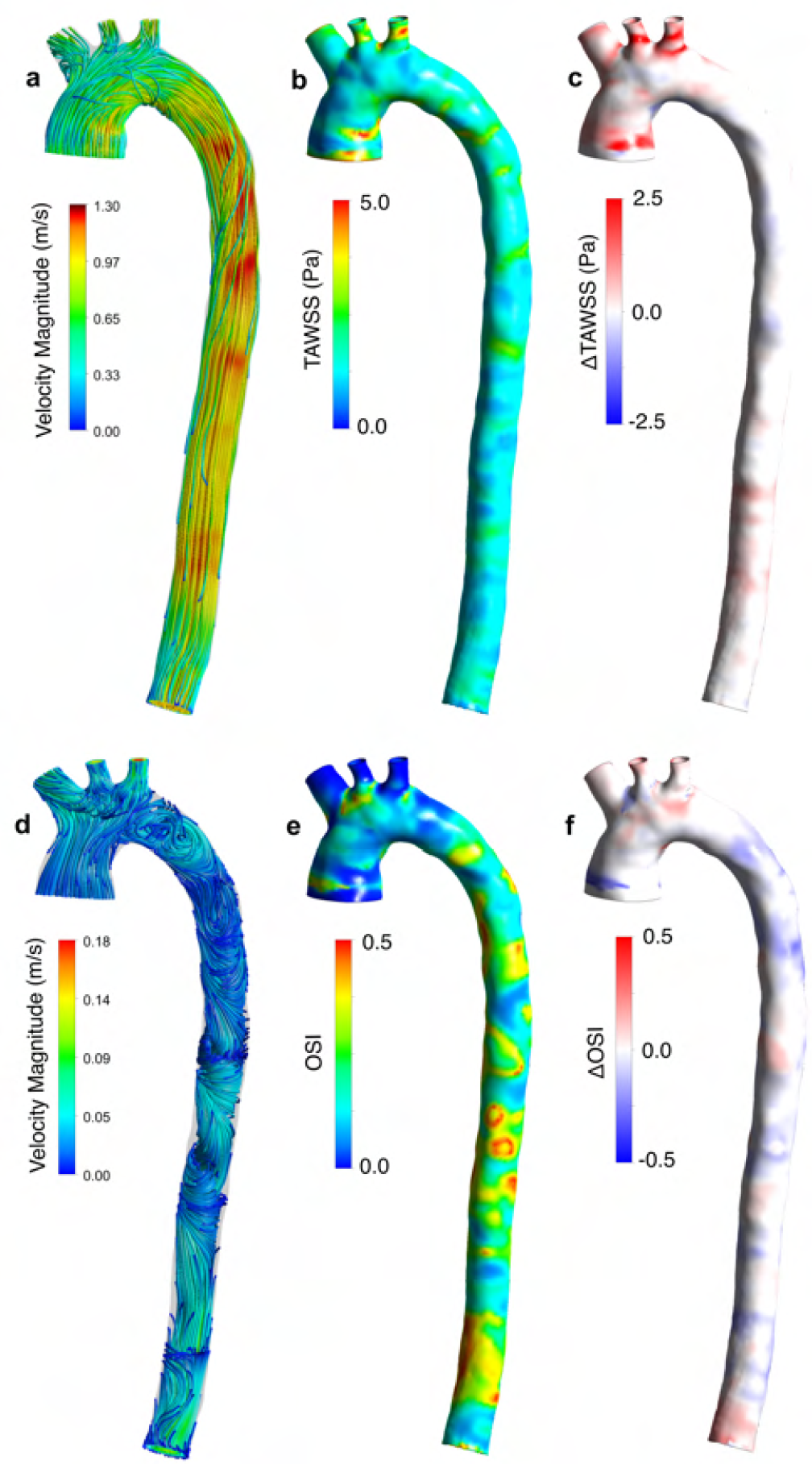
a) Velocity streamlines at peak systole, b) TAWSS contours from the CS, clipped at 5 Pa, c) contours of TAWSS difference between CS and RS, d) velocity streamlines during diastole, e) OSI contours from the CS f) contours of OSI difference between CS and RS.

High TAWSS values (above 5 Pa, according to Lantz et al. (2012); Peng et al. (2019)) are observed in the ascending aorta near the inlet, within LCC and LSA, and at their bifurcation, in agreement with other studies (Boccadifuoco et al., 2018; Lantz et al., 2011). These regions are exposed to high-velocity flow near the wall (Figure 6a). Regions of high OSI are observed in the DA where disturbed flow develops during diastole, and the WSS vector becomes highly misaligned with its average (Figure 6d). OSI was low within the supra-aortic branches, though isolated regions of high OSI are seen at their bifurcations and in the ascending aorta. This effect has been attributed to low backflow in the branches and the pulsatile separation and recirculation at their bifurcations (Lantz et al., 2011).

Although the distribution of TAWSS and OSI were qualitatively similar in both RS and CS, the rigid case exhibited substantially higher TAWSS with a 13% higher mean, a 16.4% higher maximum, and a 19.3% higher minimum value than the compliant case. These differences are concentrated in the proximal aorta (Figure 6c). As WSS indices have been identified as markers for disease progression, this has implications for the prognostic value of rigid-wall simulations that will be further discussed in Section 4.

## 4. Discussion

We have presented an efficient methodology to model patient-specific vessel compliance in CFD simulations, adapting an existing MBM workflow (Bonfanti et al., 2017) to rely solely on MRI data as input. Using a novel MRI data fusion process to segment a healthy aorta, we validated MBM results against 4DMR for the first time. By comparing compliant and rigid simulations, we have also provided further evidence that rigid simulations may not accurately reproduce physiologically accurate velocity and WSS distributions when wall movement is significant.

In this study, simulated luminal area change matched Cine-MRI measurements closely throughout the aorta, with a mean error of less than 2% in the DA and 4.6% in the arch. In comparison, recent FSI studies observe errors of 10-20% or more using a uniform Young’s Modulus, requiring numerous costly FSI simulations to determine the most appropriate value (Boccadifuoco et al., 2018; Bäumler et al., 2020). Our method achieves much higher accuracy in wall movement throughout the aorta with substantially lower computational cost than FSI and without the need for numerous calibration runs. This efficiency would be of clear benefit in a clinical setting. Small discrepancies in the shape of the waveforms may relate to differences in pressure wave transmission, and errors in Cine-MRI data due to spatial and temporal resolution.

Simulated volume flow rate waveforms agree well with 4DMR (Fig. 2). Despite equal SV, the CS and RS slightly over- and under-predict the peak flow rate, respectively. The CS models the accumulation and subsequent ejection of flow from each aortic segment as it expands and contracts under the same pulse pressure as the RS, resulting in a narrower, higher peak. Although compliance acts to delay the peak in flow rate, it was predicted slightly earlier than 4DMR, possibly indicating an under-prediction in compliance of the aortic arch and supra-aortic branches as longitudinal compliance of the proximal aorta is not accounted for in the presented methodology. Indeed, Pagoulatou et al. (2021) observed under-predictions in proximal aortic distensibility of 20-62% when longitudinal deformation was neglected.

4DMR is known to under-predict peak velocity due to cycle averaging errors (Markl et al., 2012). Therefore, it seems counter-intuitive that velocity in the DA would be under-predicted by CFD, as we observe in Figs. 4 and 5. However, 4DMR would systematically under-predict peak velocity throughout the aorta, leading to erroneously low flow rate measurements everywhere, including the AA where we extracted our inlet flow rate waveform. Therefore, this effect would be implicitly accounted for by our inlet BC and should not contribute substantially to velocity under-prediction in CFD. As the choice of IVP has not been shown to affect peak velocity in a healthy DA, we do not believe that a 3D IVP would improve the velocity agreement in the DA (Youssefi et al., 2017). It also seems counter-intuitive that peak velocity increases along the DA when 5-10% of flow is lost to the intercostals. However, the luminal area decreases by about 20% from DA1 to DA4, leading to a progressive increase in peak velocity.

We believe that the under-prediction in velocity is predominantly caused by partial volume effects and poor signal-to-noise ratio of 4DMR in the near-wall region, which contribute to an erroneously thick boundary layer of *≈* 5 mm, exceeding the expected thickness of ≈ 1mm that CFD accurately captures (see 4). Excessive boundary layer thickness would lead to a nonphysiological concentration of flow in the centre of the lumen. Therefore, *at equal flow rate* on a given plane, we would expect CFD to yield a lower peak velocity than 4DMR as the total flow is distributed across a larger area. We indeed observe this at T1 at DA3 and DA4, where peak flow rate and area are well-matched, but peak velocity is under-predicted by the simulations. This effect may be obscured by the over-estimation in peak flow rate in the compliant proximal aorta at T1, where peak velocity agrees very closely. Peak velocity is also under-predicted throughout the aorta at T2, where simulation data was extracted to exactly match the 4DMR flow rate at each plane. In the RS, this effect is obscured by the smaller cross-sectional area, which acts to increase peak velocity. Lumen areas may also be over or under-predicted as they were calibrated with Cine-MRI measurements with an associated uncertainty of ±6% in the DA. Any error in the area would compound or mitigate the impact of the thick boundary layer in 4DMR and might lead to the varying degree of velocity under-prediction observed between planes (10-28%).

We observed a moderate qualitative agreement in velocity distribution between CFD and 4DMR. However, at the arch, flow is biased to the left (outer) side in 4DMR, which CFD did not capture, which may be improved with a 3D IVP as this has been shown to improve accuracy in the proximal aorta Pirola et al. (2018); Armour et al. (2021). In the DA, the agreement is good at T1 but less favourable at T2, where simulated high-velocity regions in the DA are similar in shape but rotated clockwise by about 90 degrees compared to 4DMR. At T2, Simulations predict a high-velocity region concentrated towards the posterior wall, while 4DMR shows a concentration on the left-side wall that rotates along the DA, indicating flow helicity. As discussed in Section 2, a 3D IVP may better predict this helical flow in the DA at T2, which we will investigate in a future study.

The presence of intercostal arteries *in-vivo* may disturb the flow near each bifurcation and affect the velocity distribution throughout the DA in a way that is not simulated. 4DMR imaging errors can be substantial and are also likely to influence the agreement between CFD and 4DMR (Montalba et al., 2018; Ebel et al., 2019; Puiseux et al., 2019; Bock et al., 2019; Demir et al., 2021; Kroeger et al., 2021; Casciaro et al., 2021; Rose et al., 2016). The impact of turbulence modelling, outlet boundary conditions, rheological assumptions, aortic morphology and movement across the cardiac cycle may also affect agreement.

Without accurate resolution of the boundary layer or the location of the wall in 4DMR, WSS indices extracted from these images will incur significant uncertainties (Miyazaki et al., 2017; Piatti et al., 2017; Zimmermann et al., 2018). Despite predicting a lower peak flow rate and velocity than the CS, the mean TAWSS in the RS was 13% higher, providing further evidence that rigid-wall simulations may substantially over-predict WSS if wall movement is significant, even under identical, patient-specific and physiological pressure and flow conditions. The influence of WSS on the onset and progression of a multitude of cardiovascular diseases has been widely demonstrated (Mazzi et al., 2020). For example, specific WSS distribution characteristics have been identified as markers of aneurysm development (Chung and Cebral, 2015), and regions of low WSS have been associated with numerous adverse effects, including endothelial dysfunction and the formation of atherosclerotic regions (Wee et al., 2018). Due to the strong prognostic value of WSS, our results highlight the importance of compliance modelling.

There are some limitations to the presented method. First, the MBM cannot handle large deformations due to an associated deterioration of mesh quality. Wall movement can be in the order of 10mm in pathologies such as TBAD (Bäumler et al., 2020; Yang et al., 2014), so further work aims to improve the robustness of this technique to large deformations. Second, our methodology considers wall movement in the surface-normal direction, so any longitudinal compliance or bulk movement of the aorta is not modelled. Additionally, the effects of surrounding tissues are not considered, and the wall is assumed to exhibit linear elastic behaviour. These assumptions may contribute to differences in velocity contours between 4DMR and CFD in the proximal aorta, where the movement of the aorta away from its diastolic centreline is most significant. These effects may also impact pressure wave transmission and contribute to the minor discrepancies in flow rate and luminal cross-sectional area curves between CFD and MRI. The limitations of the uniform IVP have been discussed in detail and will be investigated in further work. Finally, the MRI data used to inform and validate the simulations is subject to various errors, as discussed. Without high-resolution experimental data (e.g. Particle Image Velocimetry), we cannot provide a systematic evaluation of these imaging errors.

This study represents a significant methodological advance in its ability to accurately reconstruct patient-specific compliant aortic haemodynamics cost-effectively using MRI data alone. Our method exhibits numerous advantages over rigid-wall simulations and FSI simulations and could facilitate improved patient safety whilst minimising healthcare resources and reducing clinical decision-making timescales. Furthermore, our technique could be generalised to other types of cardiovascular flows and aortic diseases whose morphological features can be captured accurately with MRI.

## Acknowledgements

This project was supported by the Wellcome/ EPSRC Centre for Interventional and Surgical Sciences (WEISS) (203145Z/16/Z), the British Heart Foundation (NH/20/1/34705), and the Department of Mechanical Engineering at University College London. The authors would also like to thank the Department of Computer Science of University College London for the high-performance computing cluster resources used to perform some of the simulations.

## Appendices

## Appendix A Prismatic Layers

In all meshes used for this study, ten prismatic layers with a first-layer thickness corresponding to a *y*+ of 1 were used to ensure that the first cell height lay within the viscous sublayer of the turbulent boundary layer. The total thickness of the prismatic layers exceeded the expected boundary layer thickness, *δ*, of 1.0 mm, estimated as 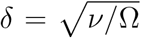, where *ν* is the kinematic viscosity, and Ω is the cycle frequency (Pier and Schmid, 2017).

## Appendix B Mesh Independence

Three successively refined meshes were used to perform a rigid-wall transient simulation with identical WK3 boundary conditions. Mesh element count approximately doubled between successive refinements, and the total prismatic layer thickness never fell below the expected boundary layer thickness of 1.0mm. Simulations were initialised using a previously converged simulation, and two further cycles were run. Less than 1% change in systolic and diastolic pressures were observed between these two cycles for each mesh. Key metrics from the final cycle of each of the three simulations are shown in Table 3.

**Table 3:**
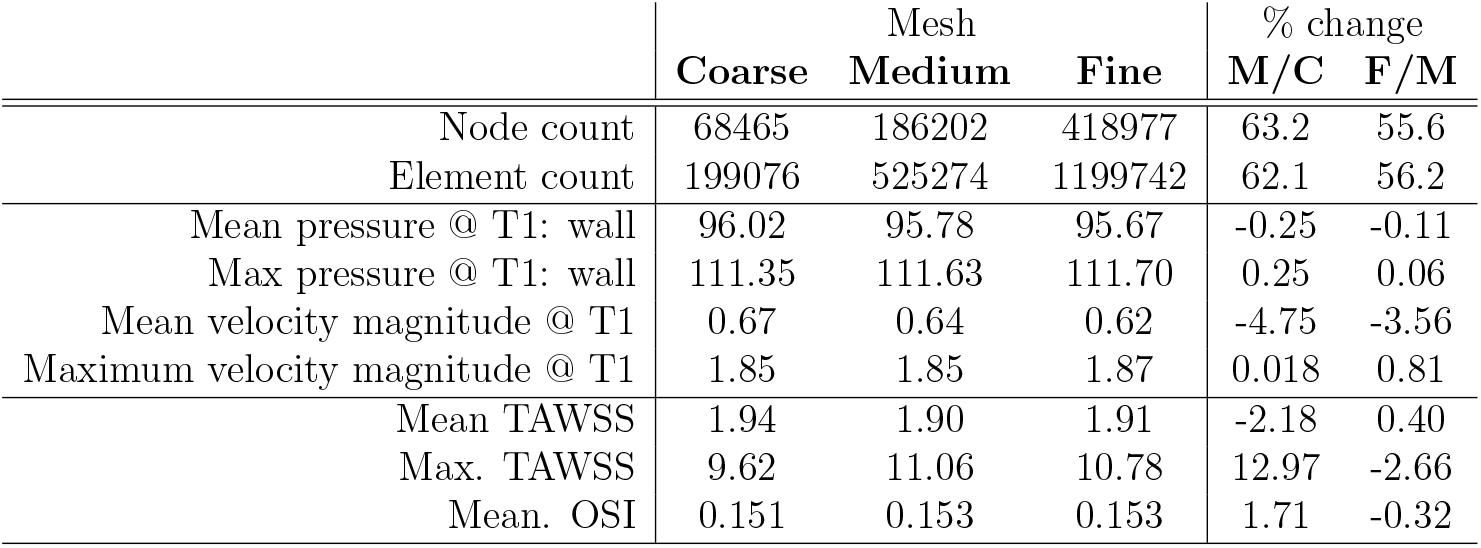
Table of key metrics from coarse, medium and fine (C, M, F) rigid-wall simulations. The final two columns show the percentage difference between medium/coarse and fine/medium meshes. T1 refers to peak systole. Velocity metrics are calculated across all cells in the domain, while pressure metrics are measured only at the wall surface.

Percentage differences between coarse and medium meshes were an average of 3.4 times higher across all metrics than between medium and fine meshes. Differences between medium and fine meshes did not exceed 3.6% for all metrics, similar to acceptable differences noted in similar studies, so the medium mesh was used for all further analysis.

## References

Armour, C.H., Guo, B., Pirola, S., Saitta, S., Liu, Y., Dong, Z., Xu, X.Y., 2021. The influence of inlet velocity profile on predicted flow in type B aortic dissection. Biomechanics and Modeling in Mechanobiology 20, 481–490.

Bäaumler, K., Vedula, V., Sailer, A.M., Seo, J., Chiu, P., Mistelbauer, G., Chan, F.P., Fischbein, M.P., Marsden, A.L., Fleischmann, D., 2020. Fluid-structure interaction simulations of patient-specific aortic dissection. Biomechanics and Modeling in Mechanobiology 19, 1607–1628.

Boccadifuoco, A., Mariotti, A., Capellini, K., Celi, S., Salvetti, M.V., 2018. Validation of numerical simulations of thoracic aorta hemodynamics: Comparison with in vivo measurements and stochastic sensitivity analysis. Cardiovascular Engineering and Technology 9, 688–706.

Bock, J., Töger, J., Bidhult, S., Markenroth Bloch, K., Arvidsson, P., Kanski, M., Arheden, H., Testud, F., Greiser, A., Heiberg, E., Carlsson, M., 2019. Validation and reproducibility of cardiovascular 4d-flow MRI from two vendors using 2 *×* 2 parallel imaging acceleration in pulsatile flow phantom and in vivo with and without respiratory gating. Acta radiol. 60, 327–337.

Bonfanti, M., Balabani, S., Alimohammadi, M., Agu, O., Homer-Vanniasinkam, S., Díaz-Zuccarini, V., 2018. A simplified method to account for wall motion in patient-specific blood flow simulations of aortic dissection: Comparison with fluid-structure interaction. Medical Engineering & Physics 58, 72–79.

Bonfanti, M., Balabani, S., Greenwood, J.P., Puppala Sapna, Homer-Vanniasinkam Shervanthi, Díaz-Zuccarini Vanessa, 2017. Computational tools for clinical support: a multi-scale compliant model for haemody-namic simulations in an aortic dissection based on multi-modal imaging data. Journal of the Royal Society Interface 14.

Bozzi, S., Morbiducci, U., Gallo, D., Ponzini, R., Rizzo, G., Bignardi, C., Passoni, G., 2017. Uncertainty propagation of phase contrast-MRI derived inlet boundary conditions in computational hemodynamics models of thoracic aorta. Computer Methods in Biomechanics and Biomedical Engineering 20, 1104–1112.

Cagney, N., Balabani, S., 2019. Influence of shear-thinning rheology on the mixing dynamics in Taylor-Couette flow. Chemical Engineering and Technology 42, 1680–1690.

Casciaro, M.E., Pascaner, A.F., Guilenea, F.N., Alcibar, J.I., Gencer, U., Soulat, G., Mousseaux, E., Craiem, D., 2021. 4D flow MRI: impact of ROI size, angulation and spatial resolution on aortic flow assessment. Physiol. Meas..

Castagna, M., Levilly, S., Paul-Gilloteaux, P., Moussaoui, S., Rousset, J.M., Bonnefoy, F., Idier, J., Serfaty, J.M., Le Touzé, D., 2021. An LDV based method to quantify the error of PC-MRI derived wall shear stress measurement. Scientific Reports 11, 4112.

Chung, B., Cebral, J.R., 2015. CFD for evaluation and treatment planning of aneurysms: review of proposed clinical uses and their challenges. Annals of Biomedical Engineering 43, 122–138.

Demir, A., Wiesemann, S., Erley, J., Schmitter, S., Trauzeddel, R.F., Pieske, B., Hansmann, J., Kelle, S., Schulz-Menger, J., 2021. Traveling volunteers: A Multi-Vendor, Multi-Center study on reproducibility and comparability of 4D flow derived aortic hemodynamics in cardiovascular magnetic resonance. J. Magn. Reson. Imaging.

Ebel, S., Dufke, J., Köhler, B., Preim, B., Rosemeier, S., Jung, B., Dähnert, I., Lurz, P., Borger, M., Grothoff, M., Gutberlet, M., 2019. Comparison of two accelerated 4d-flow sequences for aortic flow quantification. Sci. Rep. 9, 8643.

Gallo, D., De Santis, G., Negri, F., Tresoldi, D., Ponzini, R., Massai, D., Deriu, M.A., Segers, P., Verhegghe, B., Rizzo, G., Morbiducci, U., 2012. On the use of in vivo measured flow rates as boundary conditions for Image-Based hemodynamic models of the human aorta: Implications for indicators of abnormal flow. Annals of Biomedical Engineering 40, 729–741.

Gijsen, F.J.H., van de Fosse, F.N., Janssen, J.D., 1999. The influence of the non-Newtonian properties of blood on the flow in large arteries: steady F flow in a carotid bifurcation model. Journal of Biomechanics 32, 601–608.

He, F., Hua, L., Guo, T., 2021. Numerical modeling in arterial hemodynamics incorporating fluid-structure interaction and microcirculation. Theoretical Biology and Medical Modelling 18, 6.

Henningsson, M., Malik, S., Botnar, R., Castellanos, D., Hussain, T., Leiner, T., 2020. Black-blood contrast in cardiovascular mri. Journal of Magnetic Resonance Imaging, e27399.

Kousera, C.A., Wood, N.B., Seed, W.A., Torii, R., O’Regan, D., Xu, X.Y., 2013. A numerical study of aortic flow stability and comparison with in vivo flow measurements. Journal of Biomechanical Engineering 135, 011003.

Kroeger, J.R., Pavesio, F.C., Mörsdorf, R., Weiss, K., Bunck, A.C., Baeßler, B., Maintz, D., Giese, D., 2021. Velocity quantification in 44 healthy volunteers using accelerated multi-VENC 4D flow CMR. European Journal of Radiology 137, 109570.

Lantz, J., Gårdhagen, R., Karlsson, M., 2012. Quantifying turbulent wall shear stress in a subject specific human aorta using large eddy simulation. Medical Engineering & Physics 34, 1139–1148.

Lantz, J., Renner, J., Karlsson, M., 2011. Wall shear stress in a subject specific human aorta – influence of fluid-structure interaction. International Journal of Applied Mechanics 03, 759–778.

Les, A.S., Shadden, S.C., Figueroa, C.A., Park, J.M., Tedesco, M.M., Herfkens, R.J., Dalman, R.L., Taylor, C.A., 2010. Quantification of hemodynamics in abdominal aortic aneurysms during rest and exercise using magnetic resonance imaging and computational fluid dynamics. Annals of Biomedical Engineering 38, 1288–1313.

Madhavan, S., Kemmerling, E.M.C., 2018. The effect of inlet and outlet boundary conditions in image-based CFD modeling of aortic flow. BioMedical Engineering OnLine 17, 66.

Markl, M., Frydrychowicz, A., Kozerke, S., Hope, M., Wieben, O., 2012. 4D flow MRI. Journal of Magnetic Resonance Imaging 36, 1015–1036.

Mazzi, V., Gallo, D., Calò, K., Najafi, M., Khan, M.O., De Nisco, G., Steinman, D.A., Morbiducci, U., 2020. A eulerian method to analyze wall shear stress fixed points and manifolds in cardiovascular flows. Biomechanics and Modeling in Mechanobiology 19, 1403–1423.

Miyazaki, S., Itatani, K., Furusawa, T., Nishino, T., Sugiyama, M., Takehara, Y., Yasukochi, S., 2017. Validation of numerical simulation methods in aortic arch using 4D flow MRI. Heart Vessels 32, 1032–1044.

Montalba, C., Urbina, J., Sotelo, J., Andia, M.E., Tejos, C., Irarrazaval, P., Hurtado, D.E., Valverde, I., Uribe, S., 2018. Variability of 4D flow parameters when subjected to changes in MRI acquisition parameters using a realistic thoracic aortic phantom. Magn. Reson. Med. 79, 1882–1892.

Morbiducci, U., Ponzini, R., Gallo, D., Bignardi, C., Rizzo, G., 2013. Inflow boundary conditions for image-based computational hemodynamics: Impact of idealized versus measured velocity profiles in the human aorta. J. Biomech. 46, 102–109.

Pagoulatou, S.Z., Ferraro, M., Trachet, B., Bikia, V., Rovas, G., Crowe, L.A., Vallée, J.P., Adamopoulos, D., Stergiopulos, N., 2021. The effect of the elongation of the proximal aorta on the estimation of the aortic wall distensibility. Biomechanics and Modeling in Mechanobiology 20, 107–119.

Peacock, J., Jones, T., Tock, C., Lutz, R., 1998. The onset of turbulence in physiological pulsatile flow in a straight tube. Experiments in Fluids 24, 1–9.

Peng, L., Qiu, Y., Yang, Z., Yuan, D., Dai, C., Li, D., Jiang, Y., Zheng, T., 2019. Patient-specific computational hemodynamic analysis for interrupted aortic arch in an adult: Implications for aortic dissection initiation. Scientific Reports 9, 8600.

Piatti, F., Pirola, S., Bissell, M., Nesteruk, I., Sturla, F., Della Corte, A., Redaelli, A., Votta, E., 2017. Towards the improved quantification of in vivo abnormal wall shear stresses in BAV-affected patients from 4d-flow imaging: Benchmarking and application to real data. Journal of Biomechanics 50, 93–101.

Pier, B., Schmid, P., 2017. Linear and nonlinear dynamics of pulsatile channel flow. Journal of Fluid Mechanics 815, 435–480.

Pirola, S., Jarral, O.A., O’Regan, D.P., Asimakopoulos, G., Anderson, J.R., Pepper, J.R., Athanasiou, T., Xu, X.Y., 2018. Computational study of aortic hemodynamics for patients with an abnormal aortic valve: The importance of secondary flow at the ascending aorta inlet. APL Bioengineering 2, 026101.

Pons, R., Guala, A., Rodríguez-Palomares, J.F., Cajas, J.C., Dux-Santoy, L., Teixidö-Tura, G., Molins, J.J., Vázquez, M., Evangelista, A., Martorell, J., 2020. Fluid-structure interaction simulations outperform computational fluid dynamics in the description of thoracic aorta haemodynamics and in the differentiation of progressive dilation in marfan syndrome patients. Royal Society Open Science 7, 191752.

Puiseux, T., Sewonu, A., Meyrignac, O., Rousseau, H., Nicoud, F., Mendez, S., Moreno, R., 2019. Reconciling PC-MRI and CFD: An in-vitro study. NMR Biomed. 32, e4063.

Qiao, Y., Zeng, Y., Ding, Y., Fan, J., Luo, K., Zhu, T., 2019. Numerical simulation of two-phase non-newtonian blood flow with fluid-structure interaction in aortic dissection. Computer Methods in Biomechanics and Biomedical Engineering 22, 620–630.

Reymond, P., Merenda, F., Perren, F., Rüfenacht, D., Stergiopulos, N., 2009. Validation of a one-dimensional model of the systemic arterial tree. American Journal of Physiology-Heart and Circulatory Physiology 297, 208–222.

Rose, M.J., Jarvis, K., Chowdhary, V., Barker, A.J., Allen, B.D., Robinson, J.D., Markl, M., Rigsby, C.K., Schnell, S., 2016. Efficient method for volumetric assessment of peak blood flow velocity using 4D flow MRI. J. Magn. Reson. Imaging 44, 1673–1682.

Ryzhakov, P., Soudah, E., Dialami, N., 2019. Computational modeling of the fluid flow and the flexible intimal flap in type b aortic dissection via a monolithic arbitrary lagrangian/eulerian fluid-structure interaction model. International Journal for Numerical Methods in Biomedical Engineering 35, e3239.

Saitta, S., Pirola, S., Piatti, F., Votta, E., Lucherini, F., Pluchinotta, F., Carminati, M., Lombardi, M., Geppert, C., Cuomo, F., Figueroa, C.A., Xu, X.Y., Redaelli, A., 2019. Evaluation of 4D flow MRI-based non-invasive pressure assessment in aortic coarctations. Journal of Biomechanics 94, 13–21.

Tang, E., Wei, Z.A., Fogel, M.A., Veneziani, A., Yoganathan, A.P., 2020. Fluid-structure interaction simulation of an intra-atrial fontan connection. Biology 9.

Wee, I., Ong, C.W., Syn, N., Choong, A., 2018. Computational fluid dynamics and aortic dissections: Panacea or panic? Vascular and Endovascular Review 1, 27–29.

Westerhof, N., Stergiopoulos, N., Noble, M., 2010. Snapshots of Hemodynamics: An Aid for Clinical Research and Graduate Education. 2 ed., Springer US, New York, New York. pp. 191,246.

Yang, S., Li, X., Chao, B., Wu, L., Cheng, Z., Duan, Y., Wu, D., Zhan, Y., Chen, J., Liu, B., Ji, X., Nie, P., Wang, X., 2014. Abdominal aortic intimal flap motion characterization in acute aortic dissection: assessed with retrospective ECG-gated thoracoabdominal aorta dual-source CT angiography. PLoS One 9.

Youssefi, P., Gomez, A., Arthurs, C., Sharma, R., Jahangiri, M., Alberto Figueroa, C., 2017. Impact of Patient-Specific Inflow Velocity Profile on Hemodynamics of the Thoracic Aorta. Journal of Biomechanical Engineering 140.

Zimmermann, J., Demedts, D., Mirzaee, H., Ewert, P., Stern, H., Meierhofer, C., Menze, B., Hennemuth, A., 2018. Wall shear stress estimation in the aorta: Impact of wall motion, spatiotemporal resolution, and phase noise. Journal of Magnetic Resonance Imaging 48, 718–728.

